# NeuroML-DB: Sharing and characterizing data-driven neuroscience models described in NeuroML

**DOI:** 10.1101/2021.09.11.459920

**Authors:** Justas Birgiolas, Vergil Haynes, Padraig Gleeson, Richard C. Gerkin, Suzanne W. Dietrich, Sharon M. Crook

**Affiliations:** Ronin Institute; School of Mathematical and Statistical Sciences, Arizona State University; College of Health Solutions, Arizona State University; Department of Neuroscience, Physiology, and Pharmacology, University College London; School of Life Sciences, Arizona State University; School of Mathematical and Natural Sciences, Arizona State University

## Abstract

As researchers develop computational models of neural systems with increasing sophistication and scale, it is often the case that fully *de novo* model development is impractical and inefficient. Thus arises a critical need to quickly find, evaluate, re-use, and build upon models and model components developed by other researchers. We introduce the NeuroML Database (NeuroML-DB.org), which has been developed to address this need and to complement other model sharing resources. NeuroML-DB stores over 1,500 previously published models of ion channels, cells, and networks that have been translated to the modular NeuroML model description language. The database also provides reciprocal links to other neuroscience model databases (ModelDB, Open Source Brain) as well as access to the original model publications (PubMed). These links along with Neuroscience Information Framework (NIF) search functionality provide deep integration with other neuroscience community modeling resources and greatly facilitate the task of finding suitable models for reuse.

Serving as an intermediate language, NeuroML and its tooling ecosystem enable efficient translation of models to other popular simulator formats. The modular nature also enables efficient analysis of a large number of models and inspection of their properties. Search capabilities of the database, together with web-based, programmable online interfaces, allow the community of researchers to rapidly assess stored model electrophysiology, morphology, and computational complexity properties. We use these capabilities to perform a database-scale analysis of neuron and ion channel models and describe a novel tetrahedral structure formed by cell model clusters in the space of model properties and features.

**Author Summary:** Computational models of neurons and their circuits are increasingly used by neuroscience researchers as a tool to probe fundamental aspects of brain function. Here we describe a database of computational models of neurons and networks that makes it easier to evaluate and reuse these models. The models in the database are available in a standard format, called NeuroML, that makes it easier to extend and reuse the models in simulation studies using a wide range of simulation software platforms. The use of this standard format also makes it easier to characterize models in an automated way and analyze relationships across the features of simulated data from model simulations.

**Striking Image:** 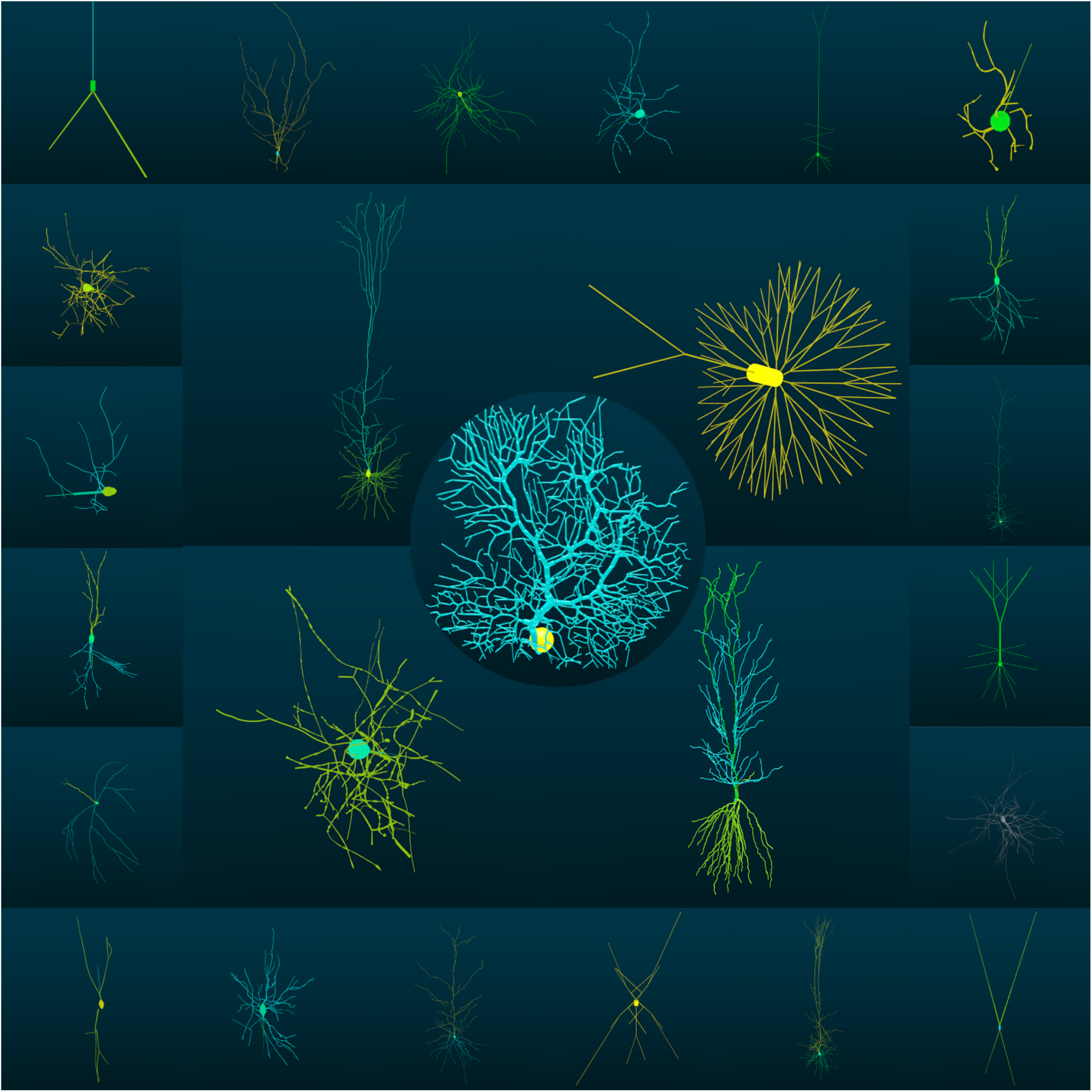

## Introduction

There are thousands of previously published data-driven neuron and neuronal network models in computational neuroscience. When a researcher wishes to create a new cell or circuit model, often existing model components could provide an efficient starting point for model development. However, the time saved with model reuse must outweigh the effort required to find, evaluate, and reproduce previous models. These tasks are supported by an ecosystem of resources for making models accessible and promoting model reuse, including model repositories and standardized model description languages. One such model description standard is NeuroML (1), which provides a modular, machine readable, simulator-independent format, and is supported by a set of software tools for describing, simulating, and analyzing models in the format ranging in scales from ion channels to large networks. We have created an online database, NeuroML-DB.org, which currently includes over 1,500 previously published models that have been translated to NeuroML. In addition to providing an interface for downloading models or their components, NeuroML-DB provides the results of systematic characterizations of the electrophysiology, morphology, and computational complexity of each model. Overall, this database adds to the existing ecosystem of resources to make it easier to find, evaluate, and reuse previously published models.

### Neuroscience data are being synthesized into increasingly complex computational models

Since the early explorations of neurons and their circuits by Golgi (2) and Cajal (3,4), humans have been fascinated by neuron diversity and complexity. In the 1950’s, Hodgkin and Huxley (5) were the first to synthesize channel electrophysiology data into a mathematical model that accurately predicted the propagation of axonal action potentials. Their approach has been extended to include a wider variety of ion channels and neuronal morphologies (6,7). More recently, data acquisition has escalated so that large data sets have been collected that describe neurons and the networks the neurons form. In parallel, exponentially increasing computing power has allowed the construction of large, biophysically-realistic network models of connected cells (8–12).

### The ability to rapidly find, evaluate, select, and reuse earlier models is becoming more important

Despite the availability of thousands of previously-created neuron models (13,14), it is still relatively difficult and tedious to find, evaluate, and select previously created models or their components like channels and synapses for reuse in a new project. Lack of consistent and easily accessible information about a model’s electrophysiology, morphology, and computational complexity makes it difficult to rapidly evaluate whether a neuron model is fit for a particular modeling purpose, resulting in the problem of *model selection*. Once a previously published model is found to be suitable, the use of heterogeneous programming and simulator languages makes it difficult to easily reuse models or model components, resulting in the problem of *model reuse*. These two problems hinder progress within the field. If current trends towards system-level modeling continue to progress toward realistic models of entire brains, the ability to rapidly leverage previously developed models will be paramount.

Online platforms for model sharing such as ModelDB (13,15) and Open Source Brain (16,17) provide rich model search and inspection capabilities to help address the model evaluation problem. Meanwhile, efforts to standardize computational neuroscience model descriptions like PyNN (18) and NeuroML (1) help address the problem of model reuse.

ModelDB is an online repository with an extensive database of published computational neuroscience models. While many models are implemented using the NEURON simulator (19,20), other simulators and programming languages are represented as well (14). ModelDB provides extensive model search and browse capabilities and, for models that are implemented in NEURON, allows users to view some aspects of the model structure like cell morphology and ion channel/synapse files. Similarly, Open Source Brain is an online collaborative environment for the development of multiscale neuroscience models. The platform leverages the structure and relationships of NeuroML models, version control provided by GitHub, and simulation visualization using the web-based Geppetto (21) platform. On Open Source Brain, model files, 3D structure, and simulation results can be inspected, and models or model components can be downloaded. Many of the models available in NeuroML-DB were originally translated to NeuroML at Open Source Brain. Similarly, an emerging online platform like Arkheia (22), where users can examine models, their parameters, and simulation results could also make it easier to evaluate models. A more narrowly focused resource, the ICGenealogy project (23,24) applies consistent and uniform stimulation protocols to ion channel models implemented in the NEURON NMODL language (25,26). The uniform protocols enable the comparison of channel models to each other, providing insight into channel model taxonomy and publication genealogy. Additionally, a new model or biological channel voltage clamp data can be uploaded to the website (24,27), which identifies similar channel models in the database allowing rapid identification of channel dynamics and a list of seed models which could be used for further fitting. The Allen Brain Atlas Cell Types Database shares models developed based on experimental data from neurons in visual cortex and the corresponding electrophysiology and morphology data (28,29); however, this resource is limited to data and models from the Allen Institute.

### NeuroML model description standardization effort promotes model reuse

The above resources make it easier to find and evaluate previously published models. Meanwhile, the standardization initiatives make it easier to reuse whole or parts of existing models. Through PyNN (18), users can specify network models composed of abstract or single compartment conductance based cell models using an expressive connectivity syntax in Python. Then by adjusting a single line of code, the model can be executed on any compatible software or hardware simulator. On the other hand, NeuroML and associated tools (30) can be used to specify network models composed of multi-compartment cells and biophysically realistic channels and synapses, as well as more abstract model formulations. Though earlier NeuroML models could only be specified using the human-readable XML format, the latest version supports compact storage using the HDF5 format (31), which facilitates the development of large, systems-level models. The modular nature of NeuroML makes it easy to extract subcomponent channel, cell, or synapse models for reuse. Additionally, these extracted components can be converted to a variety of simulator formats using automated tools (30), allowing rapid development of novel models by composition of subcomponents of earlier models.

### NeuroML database catalogs over 1,500 NeuroML models and facilitates model evaluation

Ideally, to rapidly develop a novel model, a user would be able to use an online resource to simultaneously evaluate many models and then easily select models or their components for reuse. While the model repositories described above help with locating and evaluating models, and modular and simulator-agnostic languages help with the reuse of model components, no online resource exists that combines rapid search, deep model inspection, and evaluation of features with the modular architecture of NeuroML and exposes the features via an automated interface. To make progress towards this vision, we developed the NeuroML Database and its web-based interface (32,33) (https://neuroml-db.org), catalogued published models translated to NeuroML, added model search and extensive characterization features, and implemented a web accessible web-based Application Programming Interface (API) to make it easier for researchers to evaluate, select, and reuse these models.

#### Models and search are integrated with the computational neuroscience community

NeuroML-DB contains over 1,500 models. Previously developed keyword and ontology-based search functionality (33) was supplemented with the integration of search features of ModelDB (https://senselab.med.yale.edu/ModelDB/), OpenSourceBrain.org, and the Neuroscience Information Framework (http://neuinfo.org (34). In addition to keyword search results, the ontology-based search feature can display matching cell models by their anatomical brain region locations and/or by their neurotransmitter, based on the NeuroLex ontology (35). On ModelDB, the model detail view of a published model provides links to NeuroML-DB records whenever a model appears in both databases. Similarly, using the search feature of Open Source Brain will display matching models that are also cataloged in the NeuroML database. Finally, the Neuroscience Information Framework (NIF), a federated database of neuroscience data and biomedical resources, includes NeuroML database models in its search results.

#### Standardized model characterizations are accessible online

Standardized voltage and current clamp protocols were used to characterize channel and cell models, with simulation results accessible with online interactive plots (see Results below). In addition to electrophysiological characterization, detailed cell model morphology was analyzed using L-Measure (36) and visualized, together with sample propagations of activity, using rotating online animations. Additionally, the computational complexity of cell models was assessed and compared to the reference Hodgkin-Huxley model (5) using both fixed and variable time step integration methods. Finally, the newly added models, their conversions, and their characterization data have been made available online via a machine-readable API interface.

### NeuroML database models were used to characterize the relationships across cell models

The main objective of this study was to catalog 1,500+ published cell and ion channel models within the NeuroML database, rigorously characterize them, make those characterizations available online, and describe the structure of and relationships within the cell model electrophysiology space. The electrophysiology properties of 1,222 cell models were assessed using a standardized, uniform protocol. We identified the most differentiating properties using a dimensionality reduction method and performed nested clustering analysis to identify high-density regions within the differentiating property space. This effort revealed a roughly tetrahedral structure formed by the clusters of multi-spiking cell models (see Results). We named the clusters and assessed the strength of apparent linear relationships within some of the clusters. To elucidate the underlying mechanisms of the clusters, we also characterized the contributions of ion channel currents of cell models in each cluster.

## Results

### Over 1,500 channel, synapse, cell, and network models were added to NeuroML-DB

We identified 45 publications whose models had been translated to NeuroML (see Table 1) and developed a custom, semi-automated procedure to include these models in the NeuroML database. After parsing the individual model components such as the cells, ion channels, and synapses, the database includes a total of 1,222 cell, 183 channel, 141 synapse, 27 ion concentration, and 11 network models (1,584 overall). The translation of some of these models to NeuroML was part of other efforts related to the NeuroML initiative (31,37,38). Additional models can be incorporated into the database rapidly and easily using this same procedure, and we encourage readers to refer us to any other published models that have been translated to NeuroML.

**Table 1:** The publications whose models have been translated into NeuroML and indexed by NeuroML-DB. Model counts include the top-level network or cell models as well as any distinct subcomponent synapse and channel models.

| Publication | Model Count | Short Description of Models |
| --- | --- | --- |
| Markram et. al. (2015) (8) | 1035 | Detailed cortical neuron models |
| Traub et. al. (2005) (9) | 154 | Cortical-thalamus circuit, geometric neuron models |
| Gouwens et. al. (2018) (12,39) | 106 | Detailed and point neuron models of visual cortex |
| Bezaire et. al. (2016) (10) | 42 | Hippocampus CA1, geometric neuron models |
| Traub et. al. (2003) (40) | 30 | Ion channel models |
| Prinz et. al. (2004) (41) | 28 | Crustacean stomatogastric ganglion circuit, point models |
| Maex & De Schutter (1998) (42) | 25 | Cerebellum, detailed neuron models |
| Vervaeke et. al. (2010) (43) | 21 | Cerebellum, detailed neuron models |
| Izhikevich (2003) (44) | 20 | Point neuron models with many behaviors |
| Hay et. al. (2011) (45) | 15 | Detailed cortical pyramidal neuron models |
| Smith et. al. (2013) (46) | 13 | Visual cortex, detailed neuron models |
| Dura-Bernal et. al. (2017) (47) | 10 | Motor cortex M1, point neuron models |
| Migliore et. al. (2005) (48) | 10 | Detailed hippocampus pyramidal neuron model |
| Pospischil et. al. (2008) (49) | 9 | Point neuron models of cortical and thalamic cells |
| Teeter et. al. (2018) (50) | 8 | Cortical point neuron models |
| Migliore et. al. (2014) (11) | 6 | Detailed olfactory bulb neuron models |
| Boyle & Cohen (2008) (51) | 6 | Point model of *C. elegans* muscle cell |
| Hodgkin & Huxley (1952) (5) | 4 | Point model of squid giant axon |
| De Schutter & Bower (1994) (52) | 2 | Detailed cerebellar Purkinje cell model |
| Brunel (2000) (53) | 1 | Network model of excitatory and inhibitory point neurons |
| Pinsky & Rinzel (1994) (54) | 1 | Two compartment model of CA3 neuron |
| Fitzhugh (1961) (55) | 1 | Point neuron model |
| Solinas et. al. (2007) (56) | 1 | Geometric cerebellar Golgi cell model |
| Ferguson et. al. (2013) (57) | 1 | Point model of CA1 interneuron |
| Additional ion channel models from: Korngreen & Sakmann (2000), Reuveni et. al. (1993), Poolos et. al. (2002), Migliore et. al. (2010), Kole et. al. (2006), Colbert & Pan (2002), Wang et. al. (1996), Avery & Johnston (1996), Adams et. al. (1982), Shu et. al. (2007), Magistretti & Alonso (1999), Köhler et. al. (1996), Rettig et. al. (1992), Ramaswamy et. al. (2015), Connor & Stevens (1971), McCormick & Huguenard (1992), Huguenard et. al. (1988), Hamill et. al. (1991), Astman et. al. (2006), McCormick et. al. (1993), Destexhe et. al. (1996) |  |  |

### Channel electrophysiology was characterized using ICGenealogy project voltage clamp protocols

The ICGenealogy project has developed a uniform set of voltage clamp protocols (24) that can be used to stimulate ion channel models and compare their responses (and compute their similarity indices) in an automated fashion. These protocols can be used to differentiate a variety of ion channel behaviors. While the ion channel equations stored in the NeuroML Database are stored in a structured, cross-platform XML format, it can be difficult to rapidly assess the broad ion channel behavior properties from the model equations and their parameters alone. To facilitate this assessment, we subjected the ion channels stored in the database to a subset of ICGenealogy protocols and made the model response waveforms available online in the form of interactive plots (see Figure 1). These plots allow the user to rapidly gauge the general dynamics of an ion channel model.

**Figure 1:**
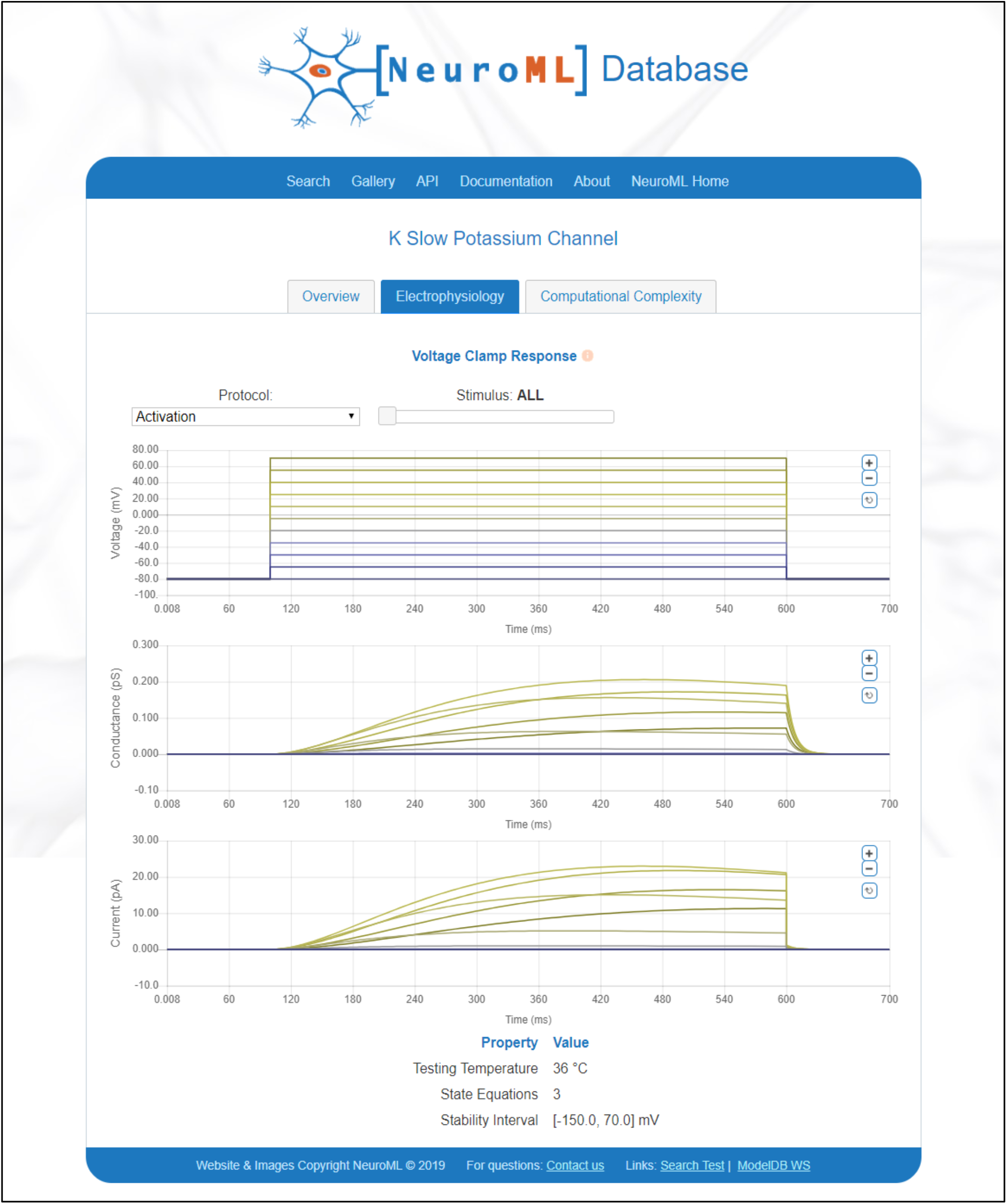
NeuroML-DB displays voltage clamp characterizations of all indexed channel models. This screenshot from NeuroML-DB depicts the characterization of an example potassium channel model (58). In addition to the activation protocol, users can view responses to deactivation and inactivation protocols using the drop-down list widget. The slider and plot zoom controls can be used to inspect individual traces. The above plots can be viewed online (59) and also accessed programmatically using the NeuroML-DB API.

For each ion channel model, the plots show the channel model input voltage and output conductance and current levels recorded over the course of simulations. A choice of protocols (Activation, Deactivation, and Inactivation) can be selected from a drop-down list and an interactive slider and plot zoom controls can be used to inspect individual traces. The waveforms are also machine accessible via the NeuroML database API. The reversal potentials and input voltage waveforms were identical to those used in ICGenealogy (23).

### Cell morphology was characterized using L-Measure and visualized using BlenderNEURON

Cell models vary in their morphological detail. Some are single-compartment “point neurons”, while others include detailed, reconstructed morphologies. Similar to channel models, while cell models are defined in a machine-readable NeuroML format, it can be difficult to visualize the 3D shape of a cell model just from the coordinates of its neurite compartments. On NeuroML-DB, we made it easy to immediately view an animated 3D rendering of each cell model with more than one compartment (see Figure 2B).

**Figure 2:**
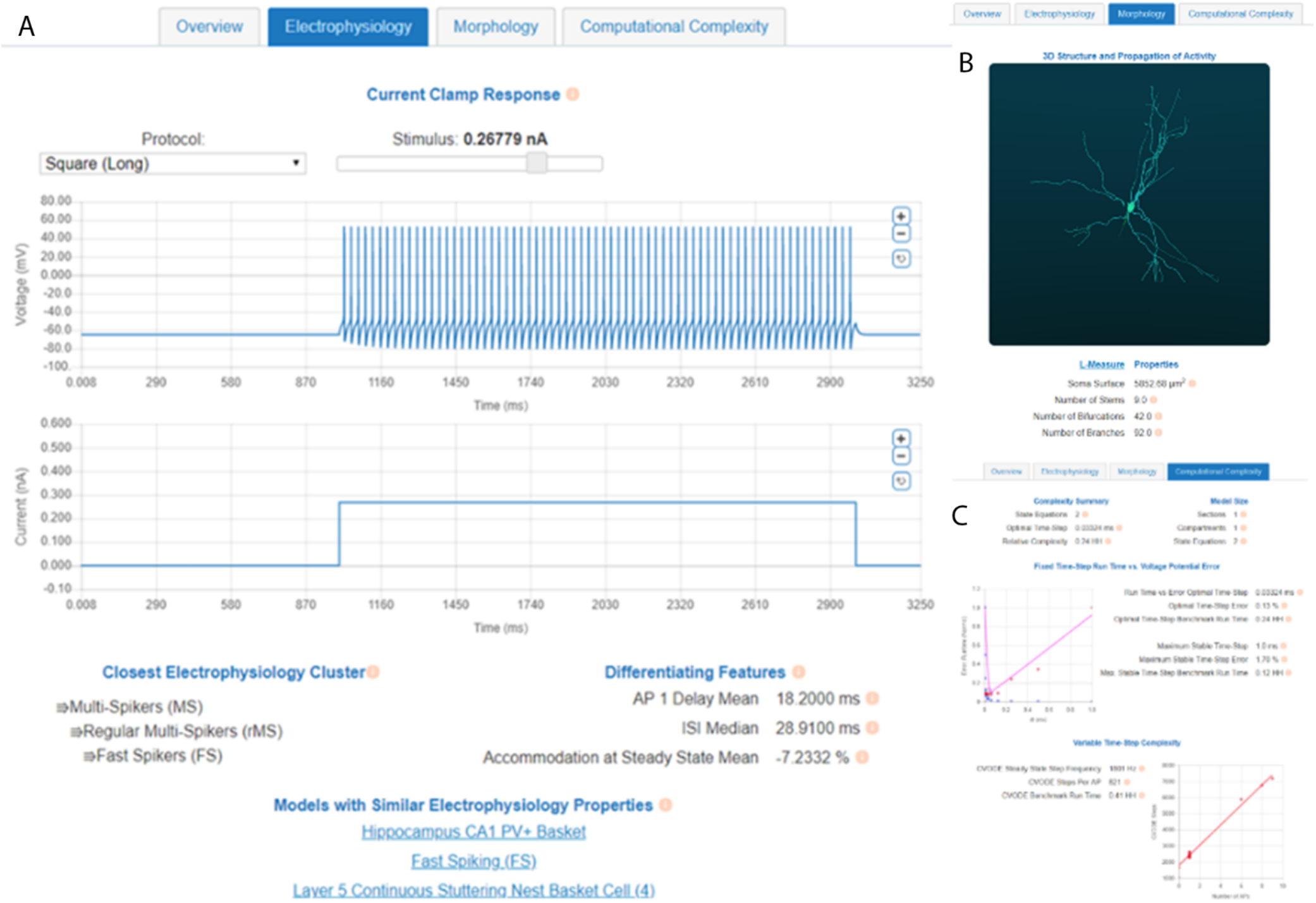
Screenshots of characterizations of a Descending Axon Cell (8) model available through the NeuroML-DB interface. A) An electrophysiology tab shows plots of current clamp responses, closest electrophysiology clusters, differentiating property values, and other cell models with similar behaviors. B) A morphology tab shows animated 3D visualizations of multi-compartment cell model geometry and electrical behavior, as well as a list of morphology metrics as computed by the L-Measure software package. C) A computational complexity tab shows model equation counts and cell model simulation speed comparisons relative to the Hodgkin-Huxley model. An example of the above plots can be viewed online (62).

To visualize it, each cell model’s morphology was first aligned along its first principal component and a 360° view of the model is shown from a slightly tilted, horizontal orbit. Additionally, a square current is injected into the model cell’s soma, and the resulting action potential propagation (if any) is shown as illuminated compartments. The animations are in the form of .GIF files, which do not require any additional software or plugins, and can be embedded, in animated form, into presentations and shared on social media platforms. To create the animations, we developed an open-source tool, BlenderNEURON (60), which creates a Python interface between the NEURON simulator and the open-source 3D modeling software Blender.

In addition to visual characterization, cell model morphology was characterized by computing the morphology metrics provided by the widely used L-Measure (36) tool. The same metrics that are visible for each reconstructed cell on NeuroMorpho.org (61) are also shown on the database website (Figure 2B bottom). The full set of computed L-Measure metrics is accessible via the NeuroML database API.

### Cell model computational requirements were characterized using numerical benchmarks

When evaluating cell models for a particular purpose, one often overlooked aspect is computational complexity – for practical purposes, how “fast” does the model run? The computational complexity of a model will affect the implementation choices needed for the optimization algorithm used for parameter fitting, as well as the overall pace of model development. In general, if all other model aspects are approximately the same, a model with a lower computational complexity likely is preferable to one with higher complexity. Computational efficiency might be especially important in model reuse scenarios where a new candidate model (for example a single component in a larger system) requires a much smaller time step than all of the other model components, making it the rate-limiting step in simulating that system.

In modeling publications, the choice of time step is often reported, but rarely rigorously justified. For example, often it is not trivial to know how sensitive a particular model is to deviations from the published time step. To facilitate such assessment and allow for automated evaluation of model simulation efficiency and stability, we developed a simple approach for characterizing the computational requirements under fixed and variable time step integration methods. Fixed step integration methods advance model state simulation by a constant time interval, while variable time step methods adjust the size of the time interval depending on how quickly the cell state changes (e.g. smaller step during an action potential, larger step when the cell has reached a steady state).

#### Cell model computational requirements under the fixed time step integration method can be measured using optimal and maximum numerically stable time step sizes

When using a fixed time step integration method, the equation count of a cell model, as well as the size of the chosen time step are the most important determinants of model run-time for a given machine and simulator combination. While changing the number of model equations might be difficult, changing the simulation time step size is not. However, for a given time step size, there is a tradeoff between model error (relative to an arbitrarily small step size) and simulation runtime: increasing the time step size will decrease model runtime but will increase model error. Thus, each model must have a time step size that will balance these two concerns. We call such time step the “optimal fixed time step size” or “optimal time step” for short.

To find each model’s optimal time step, we assessed the effect of time step size on model error and on runtime using NEURON’s default fixed step integration method. For step sizes of ∼1µs to 1ms (or each model’s maximum stable step size, if 1ms resulted in numerical overflow), we found that model error obeyed a roughly linear relationship with step size (see Figure 3A), while runtime, as expected, was proportional to the reciprocal of time step size (Figure 3B). After normalizing the error and runtime to their respective maxima, and fitting the error values to an equation of the form *b* * *dt* + *c* and runtime values to *a*/*dt*, we obtained the equation for the total cost (error and runtime, Figure 3C) of each time step size:

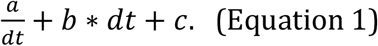

When *a, b, dt* > 0, the cost minimum, and therefore the optimal time step, occurs at

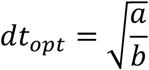

noted by the arrow in Figure 3C. To visualize how sensitive model error is to time step, we superimposed the membrane potential waveforms obtained using the smallest time step and the largest numerically stable time step (Figure 3D). To visualize how well the optimal time step captures the model behavior, we superimposed the smallest and optimal time step waveforms (Figure 3E) to find that the difference between the two waveforms was visually indistinguishable. Informally, we found that as the time step was reduced, the output waveform approached the smallest time step waveform, and the time step values near the optimal time step corresponded to a range where additional reductions in time step did not result in visually noticeable differences.

**Figure 3:**
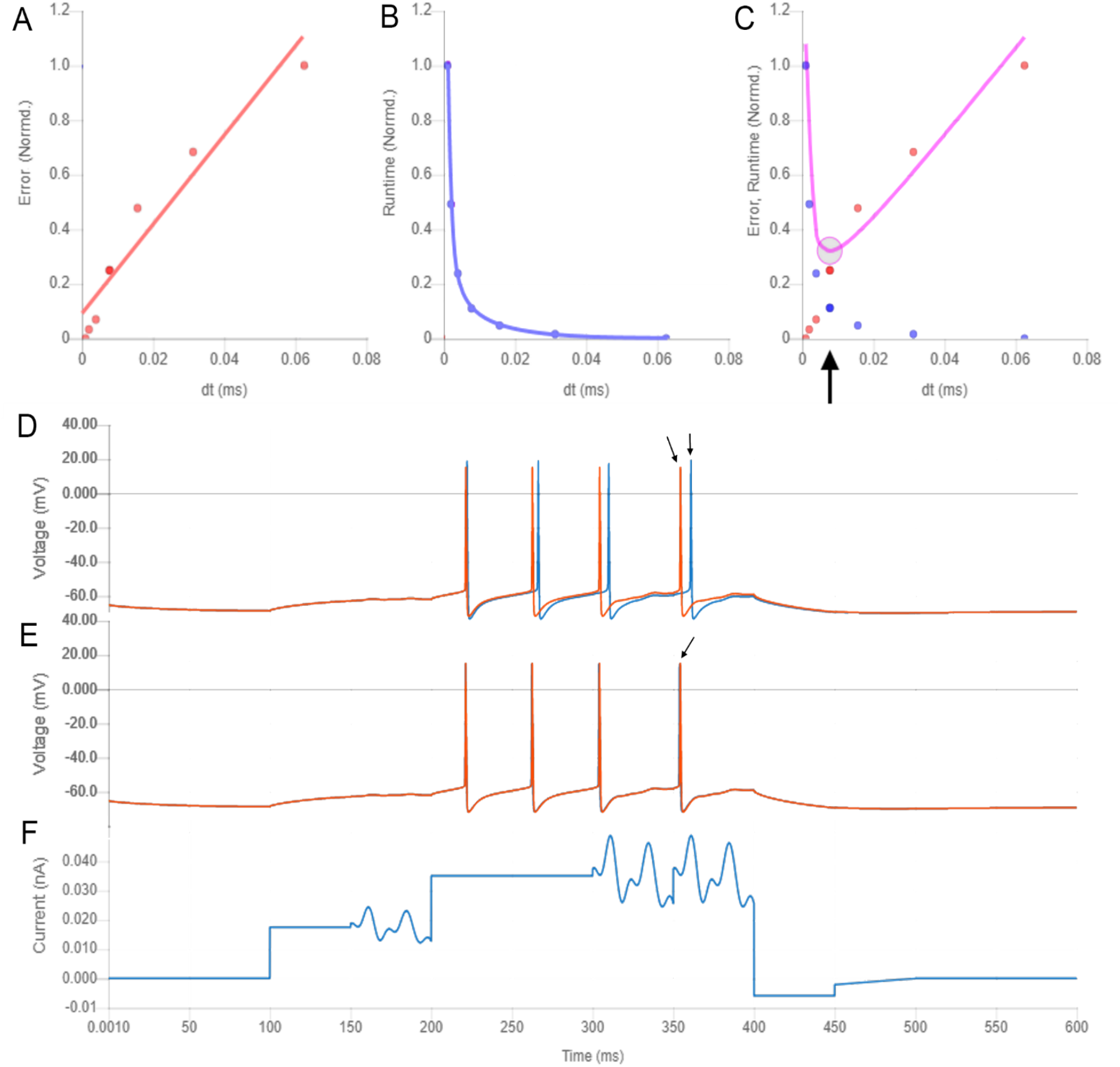
Identifying cell models’ optimal fixed time step size (model from (8)). A) Normalized point-wise model output error as a function of integration step size. Step sizes above 0.0625 ms were numerically unstable for this model. B) Normalized model runtime as a function of step size. C) The optimal time step is located at the minimum of the sum of error and runtime. D) The output responses of the model using smallest and largest stable time step (red: 1/1024 ms, blue: 1/16 ms). E) The superimposed responses when using the smallest and optimal time steps (arrow: the two traces are visually indistinguishable). F) The input waveform used to generate model responses (see Methods).

#### The computational requirements of cell models were compared to those of the Hodgkin-Huxley model

For all cell models in the NeuroML database, including the Hodgkin-Huxley model, we computed each model’s optimal fixed time step size. To assess the relative computational complexity of each model, we measured the mean time to compute a single time step of each model (based on 60 s simulations on a specific machine and simulator). Using each model’s optimal time step and the time required to execute it we determined the absolute computational complexity of each model on a specific machine and simulator combination (e.g. if the optimal time step is 0.1 ms, and one step takes 10 wall-clock ms to simulate on a given machine/simulator, then a 1000 ms simulation will take 100 wall-clock seconds to complete). To remove the machine/simulator dependence, we scaled the absolute complexities to the absolute complexity of the Hodgkin-Huxley model and obtained the relative complexities of all models in the database relative to the Hodgkin-Huxley model. This allowed us to compare each cell model’s fixed time step computational requirements, at each model’s optimal time step, to the Hodgkin-Huxley model at its optimal time step in a machine and simulator independent way (Table 2).

**Table 2:** Computational complexities (“speeds”) of select cell models in the NeuroML database. Complexities are relative to the Hodgkin & Huxley model, for which complexity is defined to be 1 HH. Top rows: models with complexities smaller than the Hodgkin & Huxley model (e.g. 0.2 HH means the model was 5 times faster than the Hodgkin & Huxley model). Bottom row: models more complex than the Hodgkin & Huxley model (e.g. 50 HH means model was 50 times slower than the Hodgkin & Huxley model). Columns: *Optimal time step*: complexity evaluation performed using NEURON fixed time step integration method, using each model’s optimal time step. *Variable*: using NEURON CVODE variable step integration method. *Maximum Stable*: performance evaluation done using each model’s maximum numerically stable fixed time step.

| Model(s) | Computational requirements<br>relative to the Hodgkin-Huxley model |  |  |
| --- | --- | --- | --- |
|  | Using<br>**Optimal**<br>Time Step | Using<br>**Variable**<br>Time Step | Using<br>**Maximum Stable**<br>Time Step |
| **Least computationally intensive models in the database:** |  |  |  |
| Teeter, et. al. (2018) GLIF model of V1 Layer 4 Spiny Cell (NMLDB ID: NMLCL001485) | 0.211 HH | 0.051 HH | 0.102 HH |
| Dura-Bernal, et. al. (2017) Izhikevich-based model of M1 PV Cell (NMLDB ID: NMLCL001664) | 0.132 HH | 0.159 HH | 0.064 HH |
| Dura-Bernal, et. al. (2017) Izhikevich-based model of M1 IT Cell (NMLDB ID: NMLCL001663) | 0.125 HH | 0.178 HH | 0.068 HH |
| Teeter, et. al. (2018) GLIF models | 0.21-0.53 HH | 0.05-0.18 HH | 0.10-0.26 HH |
| Izhikevich (2003) models | 0.17-0.27 HH | 0.29-0.77 HH | 0.10-0.14 HH |
| **Reference model:** |  |  |  |
| Hodgkin & Huxley (1952) Squid giant axon model (NMLDB ID: NMLCL001426) | 1 HH | 1 HH | 1 HH |
| Markram, et. al. (2015) models | 69-4,236 HH | 70-2,009 HH | 136-29,597 HH |
| Traub, et. al. (2005) models | 5,134-10,863 HH | 9,291-27,017 HH | 14,326-41,089 HH |
| **Most computationally intensive:** |  |  |  |
| Migliore, et. al. (2005) Multi-compartment model of CA1 pyramidal cell (NMLDB ID: NMLCL000001) | 13,652 HH | 8,588 HH | 37,706 HH |
| Traub, et. al. (2005) Multi-compartment model of LTS interneuron (NMLDB ID: NMLCL001136) | 10,863 HH | 27,017 HH | 41,089 HH |

After assessing each model’s relative computational requirements, we compared the computational complexity of the Izhikevich (44) and Generalized Leaky Integrate and Fire (GLIF) (50) classes of models to the Hodgkin-Huxley model. These classes of models are some of the simplest spiking models and are often used in large network simulations. A previous, frequently cited, computational complexity analysis (Figure 2 of (63)) based on floating point operations (FLOPS) suggested a 92-fold speed difference between the Izhikevich and the Hodgkin-Huxley model (13 vs. 1,200 FLOPS).

In our analysis, comparing empirical simulation run times, using the optimal time step method for each model, we found the actual speedup for the Izhikevich model variants (63) to be between only 3.7 and 5.7 times faster than the Hodgkin-Huxley model (Table 2). Additionally, of all models in the database that have been derived from the original Izhikevich model, the fastest model (47) had an 8-fold speedup relative to the Hodgkin-Huxley model (Table 2, top rows).

Optimal time step analysis of the simplest (1 state equation) GLIF models (50) revealed them to be between 1.9 and 4.7 times faster than the Hodgkin-Huxley model (Table 2).

It should be noted that these performance comparisons were between NeuroML models converted to NEURON using automated conversion tools. It is possible that comparing manually optimized model versions would yield different speedup ratios. While such hand-tuned model comparisons were not in scope of this research, they could be performed in follow-up studies using the methods described here.

#### Cell model complexity using variable time step size integration method can be measured using baseline steps/sec and mean steps/AP at a target firing rate

While using a fixed time step integration method is common, the popular NEURON simulator also enables simulations to use a variable time step integration method (CVODE) (64), which adjusts the size of the time step to maintain a constant local error tolerance. Because the size of integration steps varies during the simulation, evaluating the computational complexity of a cell model using CVODE requires a different method than the one using fixed time step integration. To develop such a method, we first observed the number of steps the NEURON simulator used during each millisecond to compute the response during current injections that produce an action potential (spike) (Figure 4). We noted that each model tended to have a baseline number of steps, which increased significantly during action potentials. When higher intensity current injections produced more action potentials, the total number of steps to compute 1s long simulation increased roughly linearly with the number of action potentials, across a variety of cell models. Thus, we modeled the number of steps required to compute 1 s of a cell model’s simulation as a linear function in the form of: *steps*_*sec*_ = *steps*_*base*_ + *steps*_*AP*_ * *APs*. For each model, the slope and intercept terms were fitted using a linear regression of total steps vs. number of action potentials produced in response to a series of square current injections (see Methods). Each model’s absolute 1s complexity was computed using an assumed 10 Hz (10 APs/sec) rate, and scaled to the absolute 1s complexity of the Hodgkin-Huxley model.

**Figure 4:**
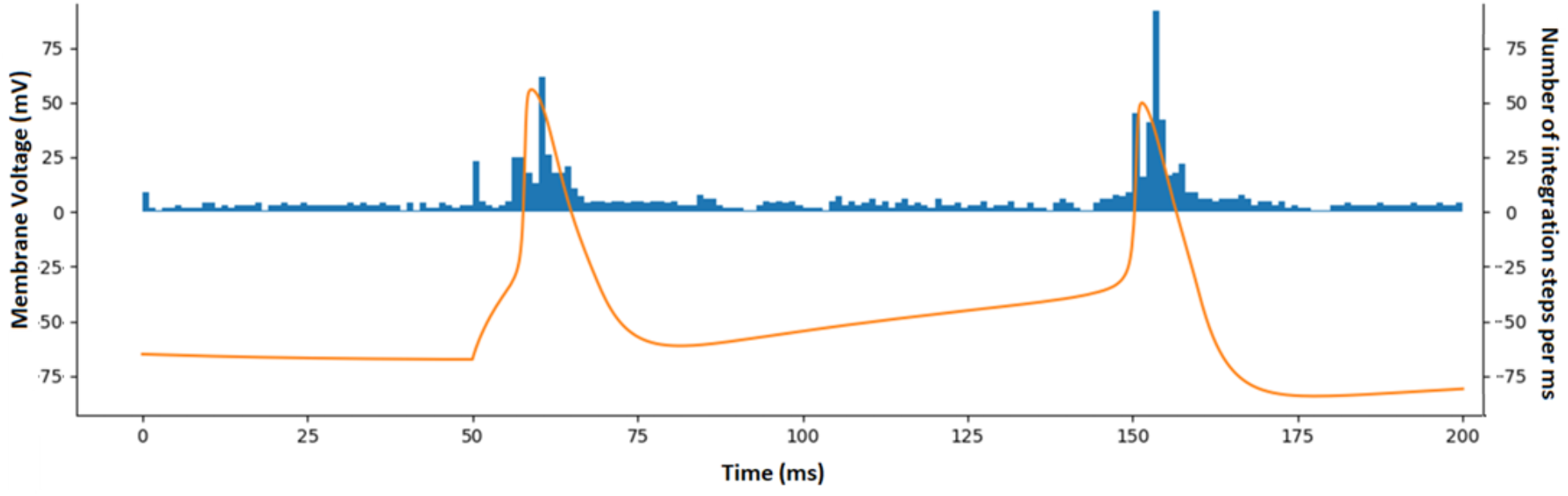
Number of integration steps required when using a variable time step integration method (NEURON CVODE) depends on the baseline step rate and additional steps per action potential (AP) for an example model (11). The plot shows membrane potential (orange) and the number of integration steps per ms computed by the simulator (blue) in response to a square current injection with onset at 50 ms. This “baseline + APs” pattern was similar across other types of input current waveforms.

As we did for the fixed step complexity measure above, we compared the GLIF and Izhikevich classes of models in our database to the Hodgkin-Huxley model (Table 2). Using variable time step method, the GLIF models were between 5.5 and 20 times faster than the Hodgkin-Huxley model, while the Izhikevich models were between 1.3 to 3.5 times faster than the Hodgkin-Huxley model.

After computing the fixed and variable time step computational complexity metrics for each cell model, we made the analyses viewable online (Figure 2C).

### Cell model electrophysiology was characterized using Allen Brain Atlas protocols and Human Brain Project electrophysiology properties

When assessing cell model electrophysiology, detailed cell model traces in response to different current clamp stimulation protocols are usually not available in the original publications. One must generally download the full model and perform the current clamp experiments manually. While this is feasible when evaluating a handful of models, it quickly becomes impractical for a large number of models. This limits the number of evaluated models, possibly resulting in a sub-optimal set of initial candidates. To facilitate this task, similarly to channel models, we have made the responses of current clamp responses to a set of standard protocols available online as interactive plots (Figure 2A). We characterized cell model responses using the electrophysiology protocols used by the Allen Cell Type Database (29), which included square, long square, pink noise, ramp, short square, and short square triple protocols (see (65) pages 7 and 15 for protocol details).

Additionally, we computed 38 cell model membrane properties described in Druckmann, et. al. (2013) (66), which were used in cell type classification by the Human Brain Project (8). Broadly, these measures assessed the properties of individual and trains of action potentials. Example action potential properties included amplitude, width, and after hyperpolarization potential, while example action potential train properties included action potential delays, inter-spike interval statistics, and degrees of spike accommodation (for full list of measures and their computation details see Table 1 and Supplementary Methods of (66), respectively). These neuron model characterizations are available as online tables (Figure 2A). Furthermore, to facilitate reuse of these properties, we implemented them as standardized tests within the SciUnit/NeuronUnit framework (67,68).

### Cluster analysis revealed the structure of model neuron electrophysiology property space

After characterizing the electrophysiology properties of cell models in the NeuroML database, we wanted to explore the structure of the space formed by the electrophysiology measures. We were interested in identifying the features that could be used to summarize cell model behavior and identify any high-density clusters (e.g. cell model types) that existed within this space. These features and cluster memberships could then be displayed online, facilitating the model selection task.

In previous work, the nomenclature established during the Petilla convention (69) provided a broad classification scheme of interneuron electrical behavior based on the consensus of the convention attendees. Later, Druckmann and others (66), used a set of 38 action potential and spike train measures of rat cortical interneurons to perform automated classification of interneurons into electrical types, which were then utilized in Human Brain Project cortical column simulations (8). Similarly, a taxonomy of mouse cortical cells performed at the Allen Institute (70), and available to explore online (71), was constructed using automated clustering of single-cell RNA sequencing data.

We ask wheter these classifications are recapitulated in neuron models. For all neuron models in NeuroML-DB, we computed the 38 properties described in (66). We reduced the dimensionality of the features by using principal component analysis (72) and used k-means and HDBSCAN clustering algorithms (73,74) to identify high-density regions of cell models within the reduced space. The analysis resulted in three levels of nested model clusters, with four clusters at the top level, two clusters in the second level, and six clusters in the third level (Figure 5).

**Figure 5:**
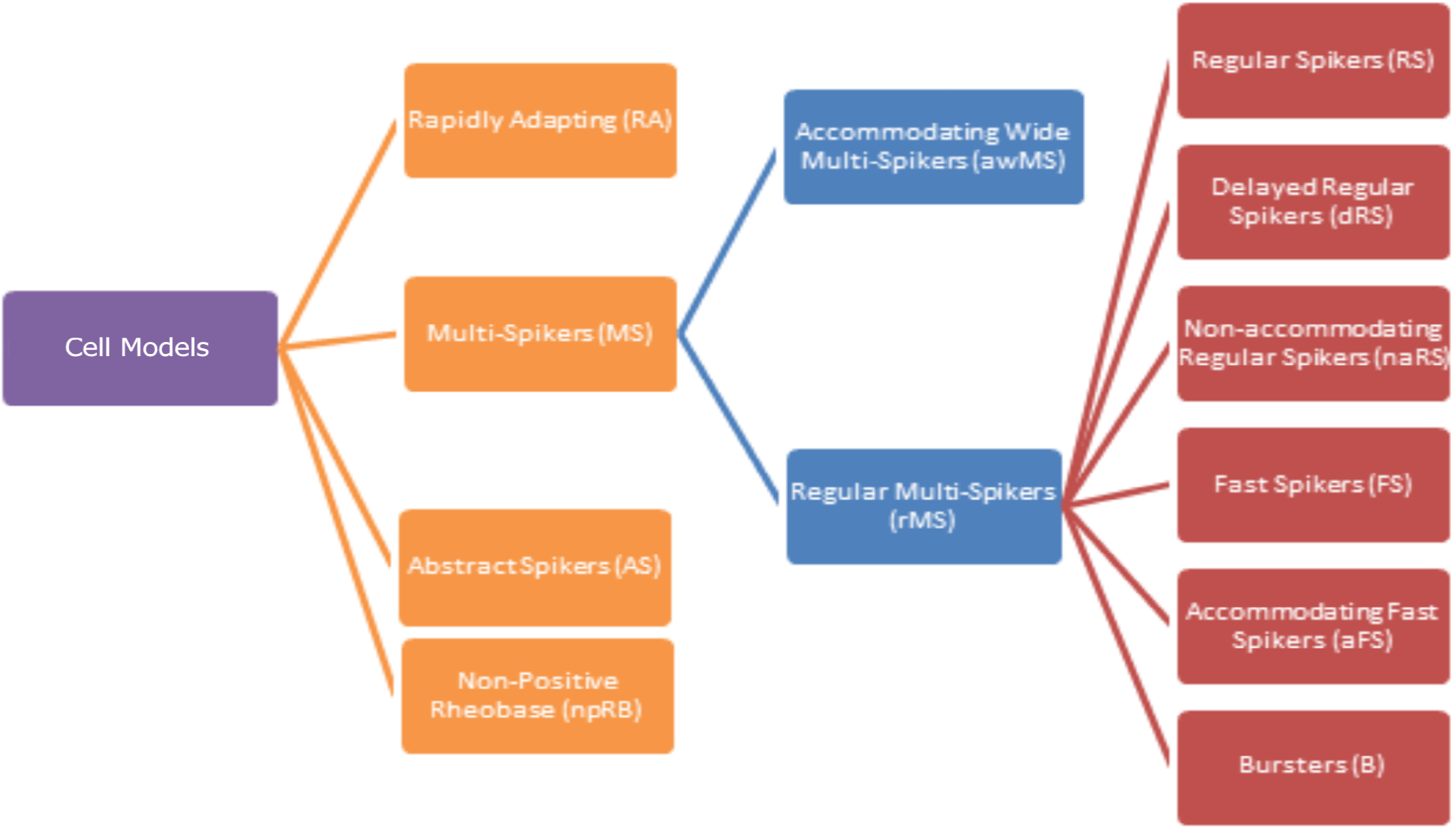
The three-level hierarchy of cell model types identified during clustering analysis of NeuroML-DB neuron models according to electrophysiology features.

At the first level of this analysis, we identified four clusters of cell models: Multi-Spikers (MS), Rapidly Adapting (RA), Abstract Spikers (AS), and Non-Positive Rheobase (npRB) (see Figure 6 for examples of models belonging to these clusters). Multi-Spikers are a large group of cell models that produce multiple spikes in response to square current injections. The Rapidly Adapting cluster contains neuron models which produce single spikes in response to square current injections. While some action potential properties can be computed for these cell models (e.g. delay to first spike, amplitude, half-width), spike train properties generally cannot be computed. Abstract Spikers are a group of neuron models for which traditional measures of action potentials do not apply, for example the GLIF cell models (50) have a resting potential at 0 mV and do not define action potential amplitude or width (Figure 6). However, the defining properties of the final six clusters in the third level are predominantly related to spike train patterns (Figure 7). For many applications, spike train patterns are of exclusive interest. While not associated with the final clusters in this analysis, the Abstract Spikers would likely belong to one of the six cluster types. Finally, the Non-positive Rheobase cluster contains neuron models that do not have a positive rheobase current and produce spikes without stimulation. Because the majority of the Druckmann properties assume positive rheobase currents, their values could not be computed (see Discussion).

**Figure 6:**
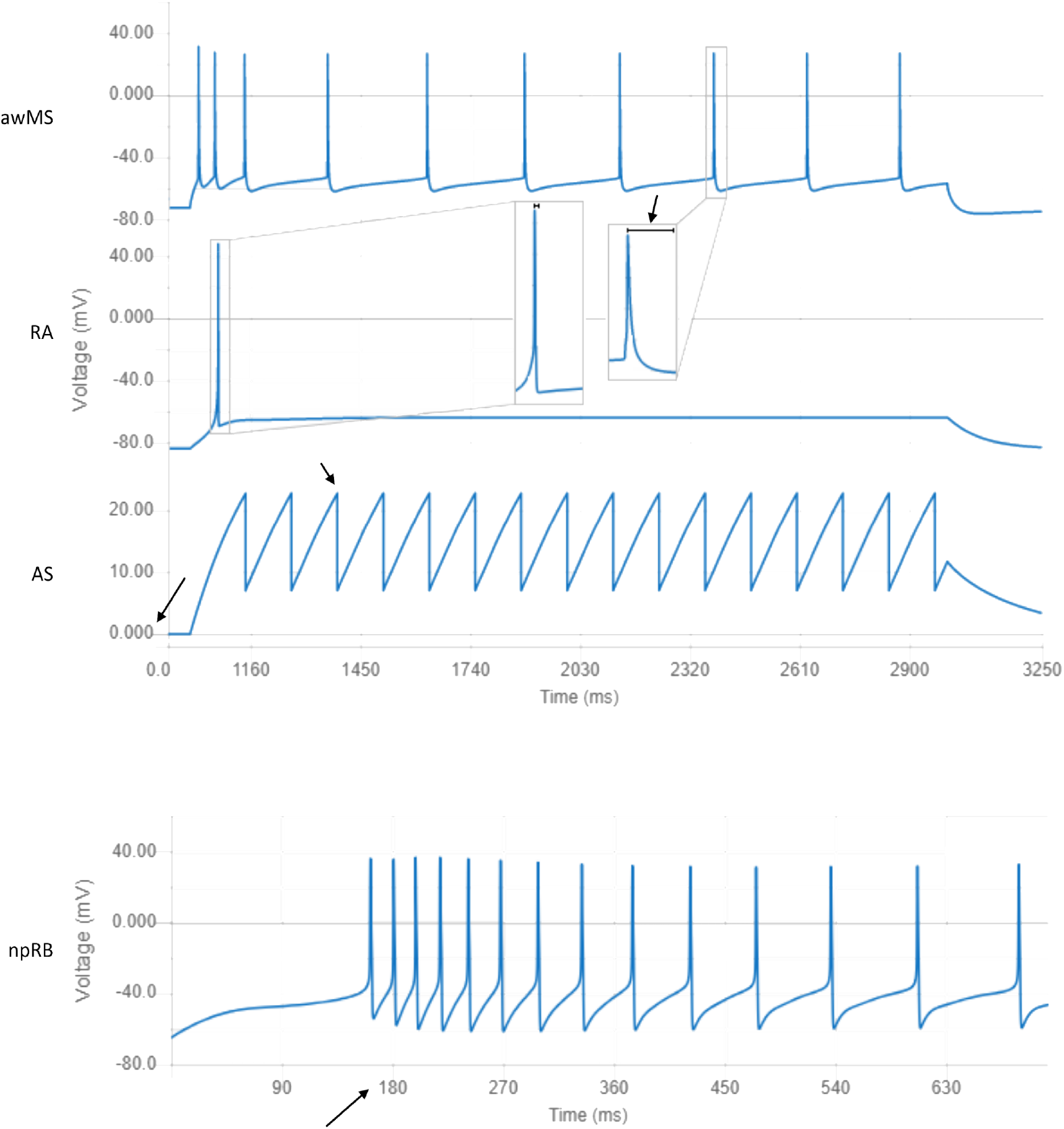
Responses of neuron models closest to the centers of clusters in the first and second levels of the cluster hierarchy. Model output examples shown above come from models referenced for each type below. awMS: Accommodating Wide Multi-Spikers exhibit spike rate accommodation and long-lasting action potentials (NeuroML DB ID: NMLCL000670). RA: Rapidly Adapting models produce single spikes in response to strong square current injections (ID: NMLCL001126). AS: Abstract Spikers do not have physiologically realistic waveforms (e.g. resting voltage at 0 mV or 0 mV action potential amplitudes. ID: NMLCL001491). npRB: Non-positive Rheobase models produce action potentials at rest (ID: NMLCL001588).

**Figure 7:**
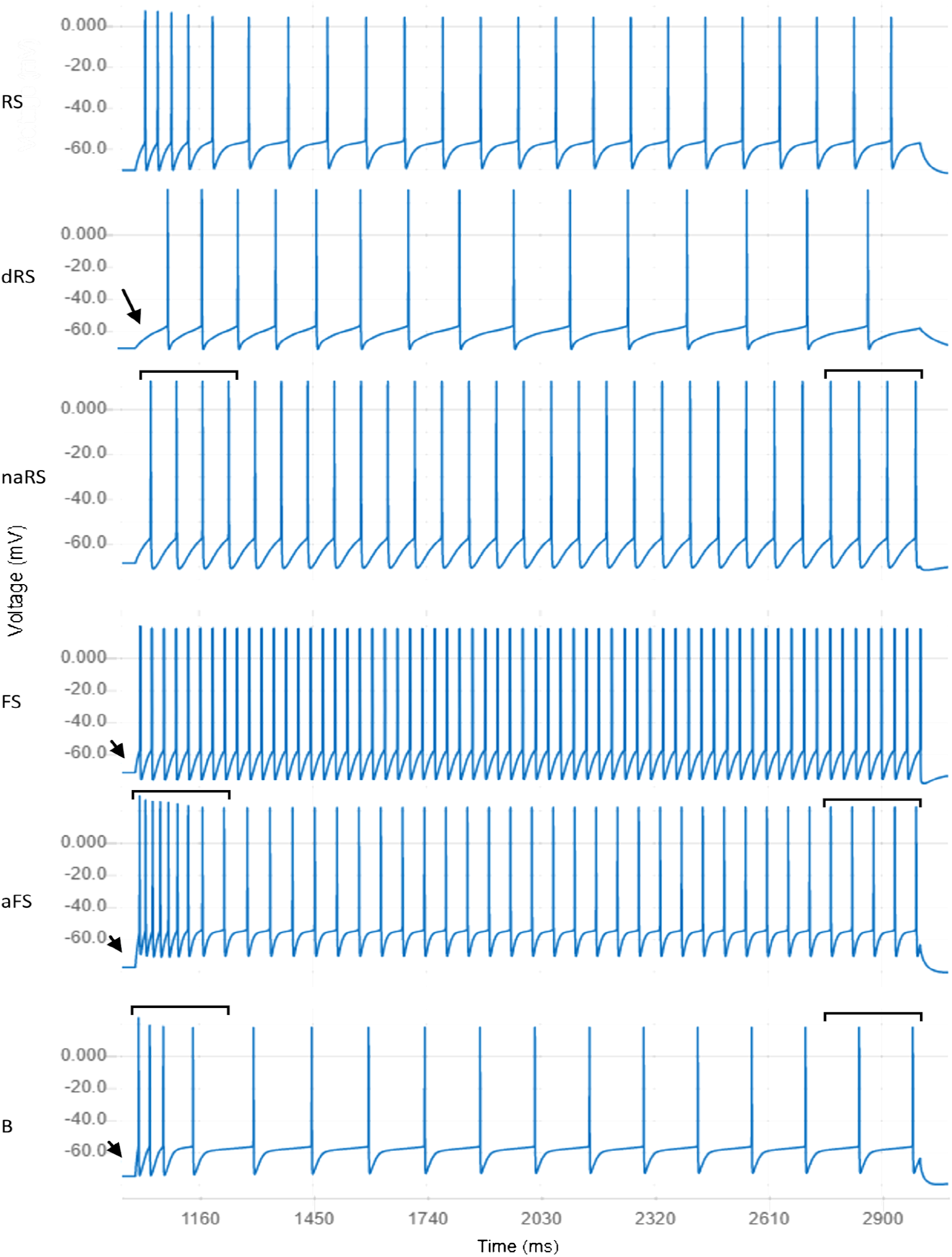
Responses of neuron models closest to centers of clusters for electrical types located in the third level of the cluster hierarchy. RS: Regular Spikers exhibit mild spike delay, some spike rate accommodation, and an average steady state spiking rate (NeuroML DB ID: NMLCL000314). dRS: Delayed Regular Spikers are similar to Regular Spikers with longer spike onset delay (ID: NMLCL000468). naRS: Non-accommodating Regular Spikers are similar to Regular Spikers but exhibit little accommodation (ID: NMLCL000829). FS: Fast Spikers exhibit rapid spike onset, little accommodation, and a high steady state spike rate (ID: NMLCL001025). aFS: Accommodating Fast Spikers are similar to Fast Spikers but exhibit some accommodation (ID: NMLCL000728). B: Bursters exhibit rapid spike onset, high accommodation, and a relatively low steady state spike rate (ID: NMLCL000203).

We then performed the same PCA and clustering analysis on the property values of the neuron models that belong to the Multi-Spiker cluster. At this second level of analysis (Figure 7), we identified a large cluster of Regular Multi-Spikers (rMS) and a smaller cluster of Accommodating Wide Multi-Spikers (awMS). The Accommodating Wide Multi-Spikers are different from the Regular Multi-Spikers in that they display much longer peak-to-trough action potential widths and exhibit spike accommodation (Figure 8, top).

**Figure 8:**
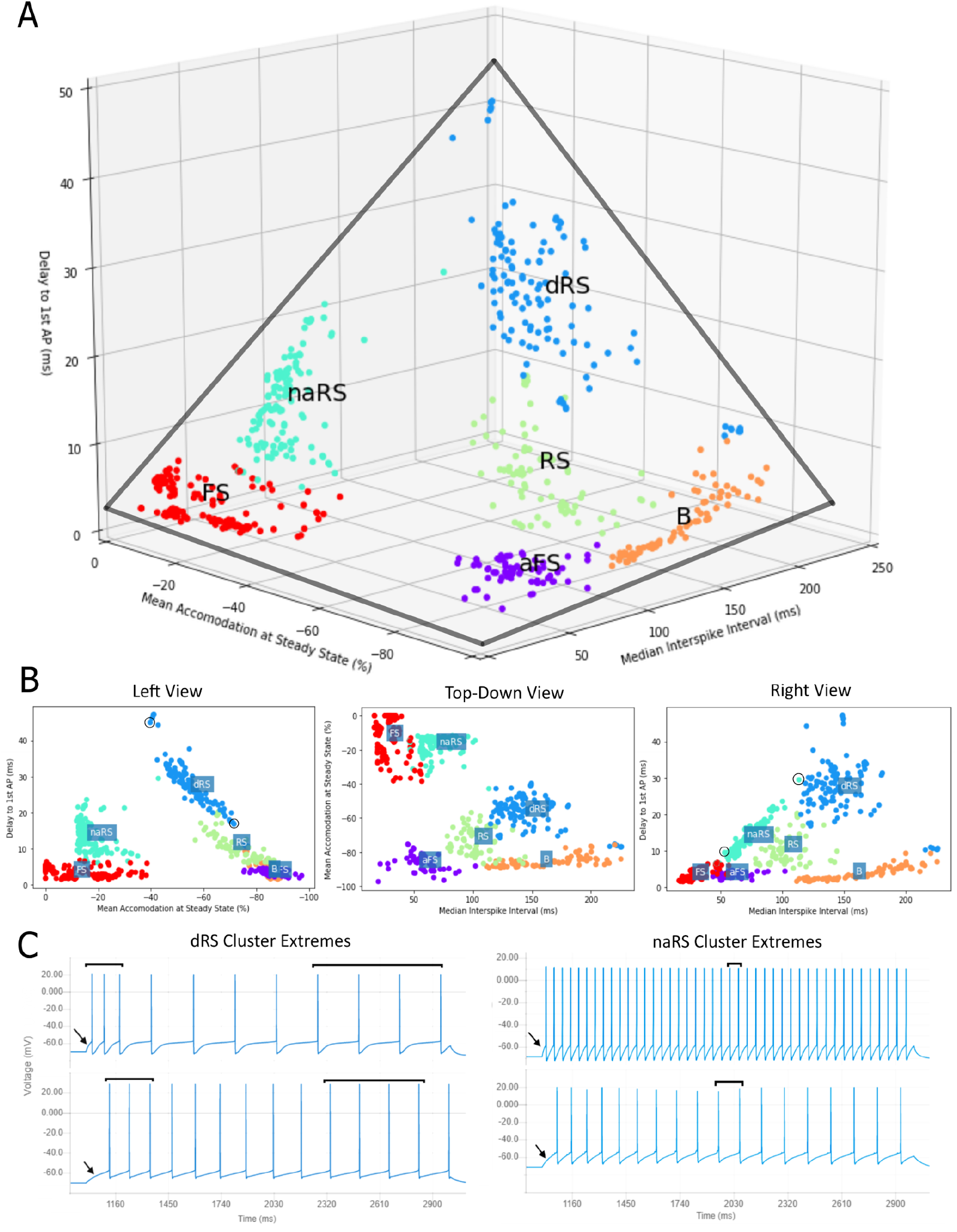
Tetrahedron of the models assigned to the high-density cluster cores of Regular Multi-Spikers. A) The level three cell model clusters plotted within the space of the three most informative properties (Delay, Accommodation, Median ISI) form a tetrahedral structure (gray lines). B) Left, Top-Down, and Right views of the tetrahedral structure. Note the strong linear relationships within the dRS and RS (delay vs. accommodation), and naRS clusters (delay vs. median ISI). C) Left: the responses of two extreme models of the dRS cluster (circled in B Left View). Note the relationship between spike delay and accommodation. Right: Extreme models of the naRS cluster (circled in B Right View). Note the relationship between median ISI and delay. For clarity, several scattered cell models that were not in the vicinity of the cluster cores are not shown (they were included in analysis, see Methods. Interactive plots that include all models can be viewed at https://tabsoft.co/32nHnNd).

Finally, we performed the same cluster identification procedure on the models that belong to the Regular Multi-Spiker cluster. At this third level, we identified 6 clusters of models: Regular Spikers (RS), Delayed Regular Spikers (dRS), Non-accommodating Regular-Spikers (naRS), Fast Spikers (FS), Accommodating Fast Spikers (aFS), and Bursters (B) (see Figure 9 for examples of each). To provide a general overview of each neuron model’s electrical behavior, we identify which cluster center is closest to the model and display the information for that cluster on the NeuroML database interface (see Figure 2A for an example).

**Figure 9.**
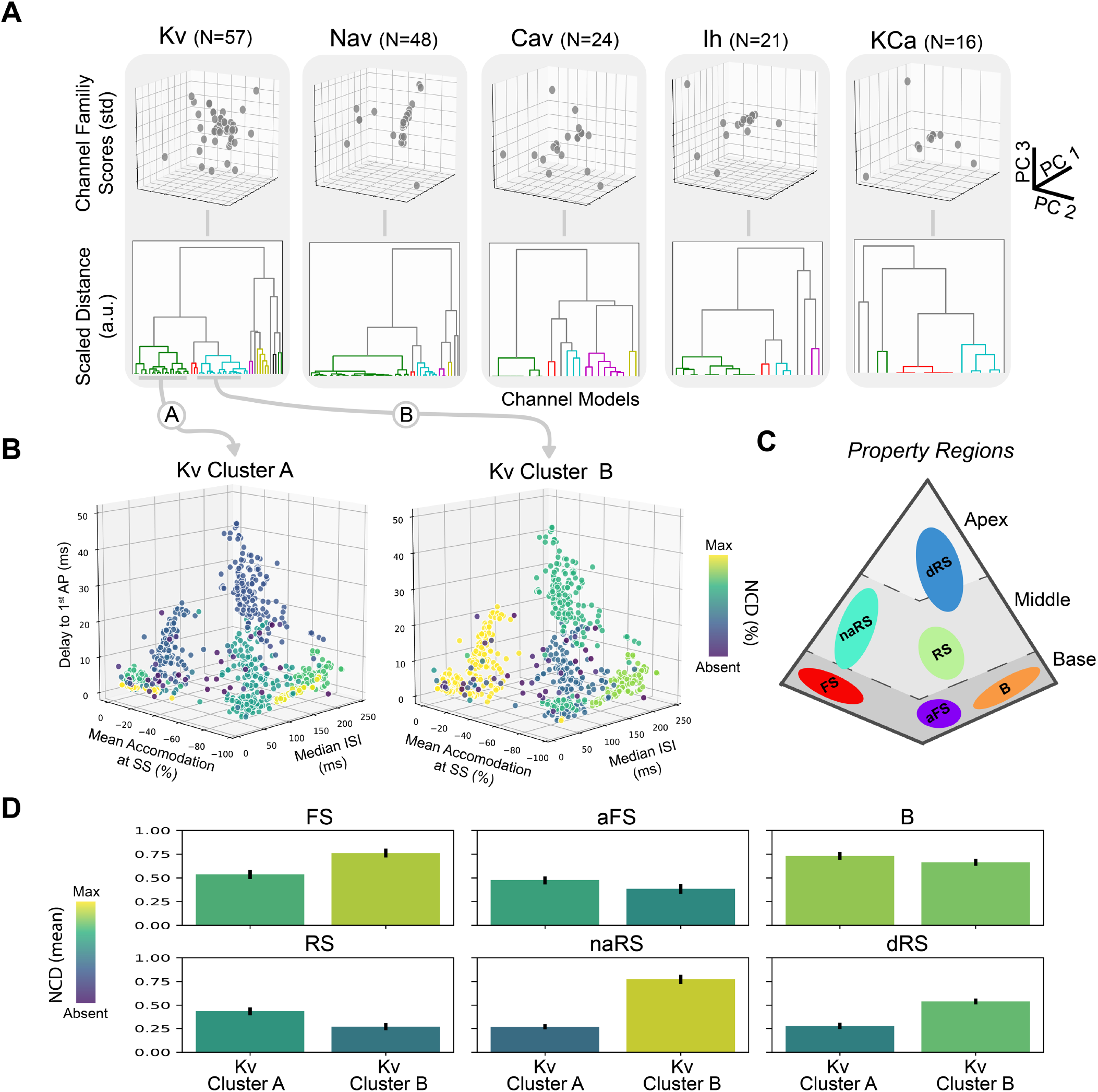
Analysis of properties of Regular Multi-Spikers based on discovered channel model sub-types. A) Dimensionality reduction and clustering reveal multiple channel model sub-types across different channel model families. B) NCDs of large potassium channel model sub-types (Z-Score<3) plotted against the electrophysiological feature space reveals global patterns for cell model parameterization. (Selected channel family clusters are indicated by gray arrows.) Note the increasing absence along the delay to 1^st^ AP axis for Kv Cluster A (Left) and the mixed patterns for Kv Cluster B (Right). C) Reference schematic of Regular Multi-Spiker high-density cluster cores in the tetrahedral property space. D) Average NCDs for each Regular Multi-Spiker cluster using all models within the base (Top), middle (Bottom-Left and Bottom-Middle), and apex (Bottom-Right) property regions (with bootstrapped 95% confidence intervals).

### Component loadings identified features responsible for clustering of Regular Multi-Spiker models

The space formed by the first few principal components of the PCA procedure is somewhat abstract and lacks an intuitive mapping to the individual features that are analyzed. To gain a more intuitive understanding of what the components represent for Regular Multi-Spikers, we first selected the first three principal components and identified single features with the highest absolute principal component weights. For the first component, the delay to first action potential had the highest weight, the second component was most heavily weighted by the median inter-spike interval, and the third component was most heavily weighted by the mean steady state accommodation percent.

When the values of these three features are plotted for the models in the cluster of Regular Multi-Spikers, we observe a roughly tetrahedral shape (Figure 8). The Fast-Spikers form the left base corner. Then, with an increased degree of accommodation, the Accommodating Fast Spikers form the center base corner. From there, Bursters, with a decreased steady state spike rate, form the right base corner. The cluster of Delayed Regular Spikers lies between the Bursters and the Fast Spikers, forming the apex of the tetrahedron. The Non-Accommodating Regular Spikers and Regular Spikers form the sides of the tetrahedron.

The sides of the tetrahedron suggested that the three properties of models belonging to the Regular Spiker sub-clusters exhibit linear relationships. We found that the Delay to First AP was strongly negatively correlated to the Mean Accommodation at Steady State for models within the Delayed Regular Spikers (Figure 8B and 8C left) and Regular Spikers (Figure 10B) clusters (dRS: Pearson r=0.94, p<0.001; RS: Pearson r=0.77, p<0.001). Similarly, the Non-accommodating Regular Spikers exhibit a strong linear relationship between the Delay to First AP and Median Inter-spike Interval (naRS: Pearson r=0.96, p<0.001). Interestingly, these relationships are weak or not significant within the models belonging to the Fast Spiker, Accommodating Fast Spiker, and Burster clusters.

### The NeuroML-DB interface facilitates rapid integration of model characterizations and metadata

As illustrated above, various voltage clamp responses of ion channel models are openly available from NeuroML-DB. Here we demonstrate the utility of NeuroML-DB for facilitating analyses of neuron models across multiple scales. We ask how the transitions along the tetrahedral structure of the discovered model property space may be explainable in terms of model ion channel densities – thus bridging channel mechanisms to neuron electrophysiology. To that end, we first characterized channel model voltage-responses using the ICGenealogy analysis protocols (24). This characterization results in compact quantitative PCA-based representations of responses for each individual channel model within their respective ionic channel families (Kv, Nav, Cav, Ih, and KCA). F inally, these PCA-based representations were used to cluster channel models into channel model sub-types using agglomerative hierarchical clustering. The resulting 22 channel model sub-types were then used to probe differences between cell model clusters for 6 multiple regular spiking clusters.

The 22 resultant channel model sub-types were not easily interpretable due to a lack of correlations to recognizable computed features (e.g., mean delay or interspike intervals). However, channel model metadata is programmatically available via the NeuroML-DB API. Channel model metadata were downloaded, and channel model names were grouped by their associated sub-types. Using these names, channel models could be characterized by common channel model names. Such names reflect the intended purpose of the models, the expertise of the original modelers, and common naming conventions used in the literature.

Consistency in channel model descriptions is observable for the larger channel model sub-type clusters. The three largest clusters belonged to the Kv (N=22 and N=19) and Nav (N=32) ion channel families, while the rest of the cluster sizes ranged from 1-7 models (Figure 9A). These largest clusters were populated with 1) delayed rectifiers and slow non-inactivating potassium channels (Kv Cluster A, Figure 9B), 2) A-type and slow inactivating potassium channels (Kv Cluster B, Figure 9B), and 3) fast transient inactivating sodium channels.

### Joint large-scale analysis integrated model characterizations across multiple scales

Channel models are used to model active conductances, which are spread across the channel membrane. The conductance densities, which may be non-uniform across the cell, can be extracted from the NeuroML model’s description. Understanding the emergence of the global structure of cell model electrophysiology requires mapping the contributions of model mechanisms at these lower scales to the computed electrophysiological features at the scale of the cell model. To gain such an understanding, we integrated the channel densities for co-clustered channel models into a single normalized channel density (NCD) representing the sum presence of the channel model cluster within the cell model. This was done for each channel model cluster and enabled joint visualization of the parameter space of channel models over the property space of cell model behaviors (Figure 9B).

The tetrahedral structure of the electrophysiological property space suggests a straight-forward way of relating the underlying mechanisms shared within a cell model cluster to this global structure. By examining how changes in average NCD correlate to movements within different property regions of the tetrahedron (Figure 9C), we observed distinct trends for the two largest potassium channel clusters. These channel clusters are known to influence long timescale dynamics. Kv Cluster A consists of voltage-gated channels with non-activating outward currents with slow activation. Kv Cluster B consists of voltage-gated channels with activating outward currents on similar timescales to Kv Cluster A. While Kv Cluster A models only have one non-activating subprocess, Kv Cluster B models have two independent subprocesses that become present at different levels of depolarization. Additionally, the inactivation subprocess of Kv Cluster B models can have time constants upward of 40-50 ms providing a slowly decreasing outward current. The differences between these two channel model clusters results in competing resonance effects that can both decrease and increase cell model spike frequency, respectively.

To what extent are transitions along the tetrahedral structure explainable by the relative contribution of the two potassium channel model sub-types? First, transitions along the base corners indicate trade-offs between accommodation and interspike intervals (Figure 9C). These transitions of the average model in the dominant base clusters (FS, aFS, B) revealed first that increasing accommodation was associated with decreasing mean NCD for Kv Cluster B shifting the NCD ratio of Kv Cluster A to Kv Cluster B to greater than 1 - illustrated in moving from the FS corner to the aFS corner. This ratio also appeared to be conserved when examining the shift from the aFS corner to the B corner. The transition was also associated with increasing mean NCD for both Kv channel model sub-types (Figure 9D, top) and increasing median ISI.

Secondly, the transition into the middle property region (increasing delay to first action potential) toward the RS cluster also conserved the mean NCD ratio of Kv Cluster A to Kv Cluster B (greater than 1) but was associated with decreasing mean NCD for both Kv channel model sub-types (Figure 9D, bottom-left). However, the transition into the middle property region between the FS corner to the naRS cluster was associated with a decrease in the mean NCD of Kv Cluster A and an increase in the mean NCD of Kv Cluster B (Figure 9D, bottom-middle). This shifted the NCD ratio to be less than 1. Interestingly, the dRS cluster in the apex of the tetrahedron also exhibits a decreasing mean NCD of Kv Cluster A, an NCD ratio less than 1, and spans the range of accommodation between the two middle clusters. These results suggest that the ratio of channel densities between the two largest potassium channel sub-types influences the degree of accommodation. Finally, a decrease in the overall channel density of delayed rectifiers and slow non-inactivating potassium channels (Kv Cluster A) increases the first spike latency evident in both sides of the tetrahedral property space.

## Discussion

The components of models published in research journals or stored in online model repositories can be difficult to quickly evaluate and re-use. The NeuroML database takes advantage of the modularity of NeuroML to efficiently provide systematic and uniform assessments of model electrophysiology, morphology, and computational complexity. By providing the results of these assessments via a freely accessible, user-friendly web interface and a machine programmable API, the website facilitates the modeler’s task of model selection with the aim of increasing researcher productivity and in turn accelerating the rate of scientific discovery. The breadth of represented models and the uniformity of their characterizations is made possible by the unique modular design of NeuroML models, which enables automated assessments. Modelers who either developed their models using NeuroML initially or have translated them to NeuroML should contact us to submit their models to the NeuroML database, where automated procedures will make the model assessment results available online without any additional effort by the model authors. Depending on the complexity of the submitted model and curator availability, analysis results can become available online within a week after submission.

### NeuroML models on NeuroML-DB can be easily extended and reused

Models in the NeuroML database are stored and made available for download in the NeuroML format. Software tools that are able to take this format as input can then be used to further process these models. For example, the jNeuroML (75) and pyNeuroML (76) libraries can be used to convert the NeuroML models to simulator formats such as NEURON (64), NetPyNE (77), XPP (78), and MOOSE (79). Additionally, it is possible to convert spiking neuron network models to PyNN scripts (18), allowing for simulations across NEST (80), NEURON, and Brian (81).

NeuroML-DB provides tested, pre-converted versions of channels (‘.mod’) and cells (‘.hoc’) in formats compatible with NEURON; these can then be used in any other software tools that can take NEURON files as inputs. Furthermore, pyNeuroML has additional features which allow automated analysis of channel model dynamics beyond what is provided by NeuroML-DB. Additionally, model construction tools like neuroConstruct (82) and NetPyNE (77) allow the composition of larger models from NeuroML components. NeuroML files can be visualized using the Open Source Brain Model Explorer (16,17,38) powered by the Geppetto (21) platform.

### Novel methods quantify computational requirements and reveal speed differences among model classes

We developed a set of measures of model computational requirements, where complexity is approximated relative to the well-known Hodgkin-Huxley model. When the absolute measures for a target and the Hodgkin-Huxley model are computed on the same machine and simulator combination, the relative measure is independent of machine speed or simulation method choice. However, the measures are dependent on the choice of integration method: fixed or variable time step. For the fixed step integration method, we developed a protocol to find the optimal time step size which balances model error and runtime. For the variable step integration method, we developed a protocol to identify the baseline number of simulation steps required to compute a unit of simulation time and the number of additional steps required to compute each additional action potential.

While it is possible to estimate model complexity by examining a model’s time constants or performing an analysis of required FLOPs (63), the measures we developed here are strictly empirical and amenable to automation because they do not require an examination of each model’s equations and parameters. Using this automated method, we were able to assess the computational complexity of 1,000+ neuron models and identify the Izhikevich and GLIF models to be the fastest classes of models in the database.

The relative complexity measures we developed have some limitations. For fixed step complexity, we utilized a rheobase-scaled input current waveform that was designed to elicit common steady state, subthreshold, spiking, and recovery behaviors. This waveform was used to produce the output response at different time step sizes and assess model waveform error. It is possible the waveform is not appropriate for all cell models and that for some models the error curve (Figure 3A) may not be linear under a different stimulation waveform. Perhaps different clusters of cell models could have different, more tailored input waveforms. If shown to be the case, we could add different protocols in future versions of the database. Furthermore, model error and runtime are weighted equally in our measure (Equation 1). Different results would likely be produced if the weights are chosen differently. Similarly, the variable step measure assumes a 10 Hz spiking rate to compute the relative model complexity. It’s possible that this spiking rate is not appropriate for some cells or provides an unfair assessment. Using an expected firing rate for each cell could be a more accurate way to assess the model’s complexity. Finally, this metric ignores any increases to step counts due to simulator events external to the cell (e.g. synaptic activity in network simulations).

### Automated assessment identifies key model electrophysiology properties and behavior clusters

Using raw traces of cell model responses to a standardized set of protocols, it is possible to identify a small number of features that capture a large portion of variability in model electrical behavior. If high density regions are present within the space formed by these features, knowing to which cluster a given model is closest also provides information about the model’s behavior. To identify such features and clusters, we performed a nested PCA and clustering analysis of the cell model electrophysiology features. This analysis was consistent with previous findings that suggested that cell electrical behavior is not uniformly distributed (66), that it exhibited clusters of models with familiar spiking behaviors such as Regular and Fast spiking, and reflected some of the important features (e.g. accommodation, delay) identified during the Petilla convention (69). Importantly, it identified the delay to first spike, median inter-spike interval, and steady state spike accommodation as three individual measures which are highly informative of cell model behavior. By making the values of these three features and the cluster membership of each model available on NeuroML-DB, we have provided an efficient public summary of each cell model’s electrical behavior.

### Top level neuron model clusters reveal opportunities to improve electrophysiology protocols

The electrophysiology protocols and computed properties used here effectively assessed the electrical behavior of multi-spiker models – neuron models with negative resting potentials that produced multiple spikes in response to 1.5x and 3.0x rheobase square current injections with no spontaneous spiking. However, this protocol and its set of electrophysiology properties did not effectively characterize cell models which did not meet these criteria.

For example, cell models in the Rapidly Adapting cluster do not produce spikes at 1.5x rheobase (1.5x rheobase is sub-threshold stimulation, e.g. RA in Figure 6). For such models, properties that rely on the production of more than one spike (e.g. second spike amplitude, or median inter-spike interval) cannot be computed. Because some of these properties are highly informative within the large group of multi-spikers, the inability to compute them makes it difficult to place the rapidly adapting cell models within the multi-spiker space. Similarly, the Hodgkin-Huxley model does not produce multiple spikes at 1.5x rheobase but does at 3.0x. It is known that electrical behavior of cells can differ under “normal” vs. “strong” stimulation conditions (69). However, the definition of “normal” vs. “strong” is not clearly defined. Druckmann and others (66), as used here, defined it as 1.5x vs. 3.0x rheobase. But is there anything intrinsically special about these rheobase multiples? For example, could the 1.5x value be already too strong for some models, and the behavior at 3.0x will not be qualitatively different? A similar issue would occur if 3.0x were not strong enough to produce behavior different from the 1.5x stimulation. Ideally, these stimulation values would be based on the dynamical bifurcation structure of the cell membrane or cell model, sampling the regions of qualitatively different behaviors. An automated method which could identify such regions in a black-box manner (e.g. without knowing the governing equations) would greatly facilitate the electrophysiology assessment of both cell models and large numbers of cells.

Another example of models which do not easily lend themselves to analysis under the current protocol is the cluster of intrinsically spiking neuron models. Because these models spontaneously produce spikes, their rheobase currents are negative (e.g. some amount of *hyperpolarizing* current must be injected to reduce the number of spikes from non-zero to zero). Because the protocols used here use positive multiples of rheobase as stimulation, little information about spikes or spike trains of such models or biological cells can be gained from using the protocol. An understanding of the diversity of intrinsically spiking or bursting cells could be used to develop an automated experimental protocol for stimulation and feature extraction to assess such cells and their models.

Finally, neuron models in the Abstract Spiker cluster have unusual membrane potential properties such as positive resting potential or 0-width or undefined-amplitude action potentials (Figure 6). Given the variety of different abstract models that could be developed, it’s not clear if a single protocol could be developed that would allow placing all such cells within the same space as the more physiologically realistic neuron models. One partial solution would be to identify separate sub-spaces for evaluating model spike train properties and action potential shape properties.

### Identifying the function of Regular Multispiker model clusters requires further comparisons to experimental recordings

Model clusters from the lowest level of the hierarchy form a tetrahedron in the 3D space formed by the features 1) delay to first AP, 2) steady state accommodation, and 3) median inter-spike interval of regular multi-spikers. This shape appears due to the strong linear relationships between delay and accommodation and between delay and median inter-spike interval within Regular Spiking models (RS, dRS, and naRS, Figure 5). The overall significance of this structure is not completely clear. If confirmed with larger and more diverse datasets of recordings from cells or simulation results from associated models, it may indicate fundamental dynamical system constraints among these three properties. AP onset, accommodation, and spike rate play important roles in neural representations of the magnitude and variation of an input signal. Since many of the models were constructed to exhibit predefined types of firing patterns (e.g., continuous accommodation or burst accommodation), the identified linear subspaces that capture the diverse types of firing patterns suggest possible compatibility between different modes of neuronal output.

The clusters that are missing are also interesting. For example, there are no high-density clusters of what could be called “non-accommodating slow spikers”, or “delayed fast spikers”, or “delayed bursters” (Figure 8). It’s possible that the tetrahedron is a product of an undersampling of potential models by the database. As additional, novel models are added, this structure might disappear. Similarly, because stochastic channels are currently not supported by NeuroML, models with stochastic firing patterns could affect the results. As the capabilities of NeuroML are further developed, the effects of such stochasticity could be tested. Finally, the tetrahedron might only reflect a structure within neuron *model* space, and it might not exist within the space occupied by experimental data. This could be tested with access to a diverse database of raw electrophysiology recordings, similar to the Allen Cell Type Database but with a larger variety of represented brain regions. More speculatively, the tetrahedron might form a volume that real biological neurons occupy due to constraints on proper neuronal function. Future investigations of how neurons with electrical behavior that falls outside the tetrahedron affect network level dynamics or how neurons of various stages of disease migrate within this space could help test this hypothesis.

### Large-scale analysis of models elucidates mechanisms across multiple scales

The joint visualization of cell model electrophysiological properties and their associated aggregated channel densities in this study is unique in that it incorporated additional information shared across models, e.g., the statistical structure of model channel densities across the database. While other high-dimensional visualizations of channel model mechanisms for neuron model databases exist (83,84), this study aimed to find common channel model mechanisms across a variety of cell and channel models spanning brain regions. Our approach demonstrated the feasibility of a functional and mechanistic mapping between the (in)activation properties of channel family model subtypes and the electrophysiological properties of cell model subtypes within the NeuroML database.

The primary goal of the combined analysis is to relate cell model electrophysiology to underlying channel mechanisms. We find that the two largest potassium channel model sub-types are associated with: 1) delayed rectifiers and slow non-inactivating potassium channels, and 2) A-type and slow inactivating potassium channels.

Intuitively, we can see that the ratio of these two discovered channel model sub-types indeed can affect the degree of accommodation (relative deceleration of spiking) of their associated cell models (Figure 9B). Further, the difference in activation thresholds provides useful information for interpreting the delay to first spike under “strong” current injection conditions (3x the rheobase). Kv Cluster B models are activated under weak stimulation while Kv Cluster A are activated only under strong stimulation of the cell model. This additional activation of outward currents from Kv Cluster A is one of the dominant causes of the delay to first spike (Figure 9B).

### Future directions

The current release of the NeuroML database focuses on the characterization of ion channel and neuron model electrophysiology, neuron model morphology, and model complexity. Based on the requests of researchers, we could add additional features to the database to help make the selection and reuse of previously published models more efficient. In future releases, based on user demand, we may provide characterizations of synapse models, and similar dimensionality reduction and clustering analysis could be performed with synapse and cell morphology data. For network models, we could provide visualizations of their structure and connectivity and display traces of network outputs in response to arbitrary inputs. Another useful feature might be to identify cell models in the database that have similar responses when provided with a set of user-uploaded electrophysiology recordings from experiments.

Finally, we’ve made the analysis code available online (85), which allows other researchers to compute the electrophysiology, morphology, and complexity properties from responses of other models or even experimental data.

## Methods

### NeuroML tools and simulator

jNeuroML v0.8.3 was used to parse and convert all NeuroML models to NEURON format (.mod and .hoc). All simulations were performed using NEURON simulator v7.5 on an Intel(R) Xeon(R) CPU E3-1240 V2 @ 3.40GHz, 32 GB RAM machine running Ubuntu Linux 16.04 LTS. For fixed step simulations, NEURON cvode_active variable was set to 0, while for variable step simulations the variable was set to 1 before starting simulations.

### Fixed time step computational complexity

For each spiking cell model that was not intrinsically spiking, the following procedure was used to compute the fixed time step complexity.

#### Input

An input current waveform (see Figure 3F) consisting of 100 ms at 0 nA, 50 ms at 0.75 rheobase (RB), 50ms of a pink noise waveform at 0.75 RB, 100 ms at 1.5 RB, 50 ms of pink noise waveform at 1.5 RB, 50 ms at -0.25 RB, 50ms of pink noise at -0.25 RB, and 100 ms at 0 nA (total length 600 ms) was injected into the center of the soma section and the voltage of the compartment was recorded. The pink noise waveform can be downloaded from (85) (file labeled “dtSensitivity.pickle”).

#### Time step sensitivity

The simulation of the above protocol was repeated multiple times, each time using a different time step size. Starting with a time step of 0.0009765625 ms (1/1024 ms), the time step size was doubled until it either reached 1 ms or the simulation became numerically unstable (“blew up”). The largest time step size that did not blow up was saved as the maximum stable step size.

#### Error

The output waveform produced using the smallest time step size (1/1024 ms) was assumed to be the reference waveform and assigned error of zero. The waveforms produced by larger time steps were compared to this waveform at 1 ms intervals. Each waveform’s error was computed as the average of point-by-point differences from a reference waveform expressed as percentages of the reference waveform min-max range (code for this function can be found in (85)).

#### Optimal time step

The optimal time step (example seen in Figure 3C) was found by identifying the minimum of the sum of time step error (Figure 3A) and time step runtime (Figure 3B). Time step error depends linearly on the time step (*dt*) and was fitted to a linear function *b* * *dt* + *c*. Runtime is inversely proportional to the time step and was fitted to a function *a*/*dt*. Fitting was performed using scipy.optimize.curve_fit function (86). The sum of these two functions (*C* = *a*/*dt* + *b* * *dt* + *c*) represents the total cost of using a particular time step to perform a simulation. The minimum cost, and therefore the optimal time step occurs where the derivative of the cost curve is zero as demonstrated in Figure 3A. Code for finding the optimal time step is available (85).

#### Mean runtime per step

Due to background operating system processes, the time required to compute one model time step is variable. For this reason, the mean runtime per time step (*runtime*_*step*_) was measured for each model. Each model was simulated without any stimulation using a time step of 0.0078125 ms (1/128 ms), for approximately 60 wall-clock seconds. To get the mean runtime per time step, the actual simulation time was divided by the number of steps required to perform the simulation. To maximize accuracy, the simulations were performed one at a time, on the same machine without any other running tasks.

#### Absolute and relative complexities

Absolute complexity *Ω*_*abs*_was defined as the total wall-clock time required to run a model at its optimal time step to simulate its output in response to the 600 ms test waveform (see Figure 3F). This complexity partially depends on the computational power of the executing machine. To obtain a complexity measure that is machine-invariant, the absolute complexity of a model can be divided by the absolute complexity of a reference model, yielding a quantity which represents a complexity multiple relative to the reference model. Here, the reference model was the Hodgin-Huxley single compartment model (5), and all model complexities were expressed relative to it. A model’s absolute complexity can be computed from the length of the input waveform (600 ms), the optimal time step (*dt*_*opt*_), and the mean runtime per step (*runtime*_*step*_) using the equation *Ω*_*abs*_ = *runtime*_*step*_ * 600/*dt*_*opt*_. Each target model’s absolute complexity *Ω*_*abs*_was divided by the absolute complexity of the Hodgkin-Huxley model to obtain the target model’s complexity relative to the Hodgkin-Huxley model (*Ω*_*HH*_).

### Variable time step computational complexity

For each spiking neuron model that was not intrinsically spiking, the following procedure was used to compute the variable time step complexity. The procedure assumes a positive rheobase current, which does not exist for intrinsically spiking models.

#### Input

Square current 1s long after a 1s delay were injected into each model’s soma section. The current amplitudes were: 0 nA, the largest known sub-rheobase current, and 11 evenly spaced currents valued between the rheobase and 1.5 times the rheobase. The number of steps the CVODE integrator used to compute the output and the number of action potentials produced in response to each current waveform was recorded (code available in (85)).

#### Baseline steps and steps per spike

The number of action potentials produced in the waveforms was linearly regressed using the *curve_fit* function (86) versus the number of simulator steps required to compute the waveforms. The intercept was interpreted as the baseline number of steps required to compute 1s of simulation (*steps*_*base*_), and the slope was the mean number of additional steps required for each additional action potential (*steps*_*ap*_, also see Figure 4). The code for this computation can be viewed in (85).

#### Absolute and relative complexities

The absolute variable time step complexity of a model was defined as the wall-clock time required to simulate 1s of model output which contains 10 action potentials (target firing rate of 10 Hz). The equation for it was *Ω*_*abs*_ = V*steps*_*base*_ + *steps*_*ap*_ * 10X * *runtime*_*step*_, where 10 was the target spike rate, and *runtime*_*step*_ was the mean runtime per step described in the “Fixed time step computational complexity” section above. The absolute complexity depends, in part, on the computational power of the executing machine. Similar to the fixed time step complexity, the machine speed factor can be removed by dividing the target model’s absolute complexity by the absolute complexity of a reference model. Here, the relative variable time step complexity of a model (*Ω*_*HH*_) was computed by dividing each target model’s *Ω*_*abs*_ by the *Ω*_*abs*_of the Hodgkin-Huxley model.

### Nested cell electrophysiology dimensionality reduction and clustering

#### Properties, Missing Values, and Transformations

All 38 electrophysiology properties from (66) were defined using reusable, Python-based tests within the NeuronUnit (87) validation framework (the tests are available in (67)). Additionally, the following four properties were used in the PCA and clustering analysis. Resting Action Potential Count was computed by counting the number of action potentials produced by the cell without current stimulation. Time to First Ramp Spike was computed by measuring the number of milliseconds required for the cell model to produce an action potential while it was injected with a ramp current increasing at the rate of 1 rheobase/sec. Finally, frequency filtering response of each cell model was assessed by fitting the number of action potentials produced in response to square current triples, spaced at frequencies ranging from 29 to 143 Hz (from (65)), to bi-sigmoidal frequency response curves (“hat”), and using the fitted inflection point locations as the frequency filter pass above- and below-filter parameters (the code for this procedure can be found at (85)).

The values of some properties of some models were missing (e.g. amplitude of 2^nd^ AP when cell only spiked once). In such cases, the missing values were replaced with either minimum or maximum possible or mean values, as deemed appropriate. The code to fill the missing values can be viewed at (85).

Some property values had strongly skewed distributions and non-linear relationships with PCA components. To reduce the effect of outliers on PCA results and to achieve closer concordance with the PCA linearity assumption, such properties were transformed using the bi-symmetric log transformation (88). This transformation could scale negative values, did not exaggerate values between ±1, and resulted in higher PCA component vs. transformed property correlation r values than the more common “offset and then take the log” method for properties with large negative and positive values. The result of subsequent clustering analysis was qualitatively similar to the result obtained when using the more common log and cube root transformations. Transformation code and properties transformed can be found in (85).

#### Dimensionality Reduction

The dimensionality of the 42 features was reduced by the use of PCA. Z-scores of filled, and bi-log transformed property values were used as inputs to the PCA function implemented in scikit-learn(89). The first N components that accounted for 95% of variance were retained (starting dimensions = 42, post-PCA dimensions = 21, n = 1222).

#### Nested Clustering

K-means (89) and HDBSCAN (74) clustering algorithms (scikit-learn), were used to group the values of principal components of cell model features. HDBSCAN with minimum cluster size of 10 was used for the first two levels of cell model clusters (in Figure 5, “multi-spiker” cluster in level one and “regular multi-spiker” cluster in level two). K-means algorithm was used in level three. There, the number of selected clusters was chosen by exploring the range of 2-10 clusters with silhouette analysis (90), and picking the cluster count with the largest silhouette score (89). Across all levels, the number of clusters identified by the algorithms was validated with visual inspection by plotting the first three PCA components.

In all levels, HDBSCAN was used to identify cell models that were not in the vicinity of the high-density cluster cores. These “noise” models were excluded when using the K-means algorithm. Post-clustering, each “noise” model was assigned to the cluster whose geometric center had the smallest Euclidean distance in PCA space. The cluster assignments for all cell models were stored in NeuroML-DB.

Linear relationships observed in Figure 5 were assessed after each model was assigned to a cluster. Pearson correlation coefficients and their p-values were computed using the SciPy Python package.

The code for the above procedures can be viewed at (85).

### Channel model electrophysiology dimensionality reduction, clustering, and parameter analysis

#### Pre-Processing and Dimensionality Reduction

Unlike the cell model data, no computed features are available for channel models at Neuro-ML DB. Instead, the voltage-clamp response data, as well as corresponding metadata, were programmatically downloaded from NeuroML-DB via the API. Pre-processing the data followed a previously established pipeline (24). The final result was a condensed representation of the temporal responses of all channel model voltage-clamp responses grouped by protocol type (activation, deactivation, and inactivation) and channel family (Kv, Nav, Cav, KCa, and Ih). Dimensionality reduction was applied to this reduced representation also following the pre-established protocol from (24). In summary, PCA was independently applied to the matrix of model temporal responses for each protocol with the number of components chosen to account for 99% of the variance. This first application of PCA reduced the dimensionality of each time series for the different protocols and removed temporal correlations while maintaining the majority of variance across channel model outputs within a given protocol. Subsequently, the individual channel models were then given a score based on the fit of the different PCA representations for each model, i.e., the log-likelihood of each model sample under the different learned PCA models. This resulted in a three-dimensional score for each channel model serving as computed feature vectors from the independent PCA spaces. The matrix of *channel family scores* was then subjected to PCA (99% explained variance retained). This second application of PCA discovered the directions of maximal variance of the channel family scores across all protocols suitable for clustering.

#### Hierarchical Clustering

Agglomerative hierarchical clustering was performed on the each of the final channel family score matrices using Ward’s minimal variance linkage (91) implemented in scikit-learn (89). A dynamic tree cut algorithm (92) was used to determine cluster cut-off values for the resulting dendrogram of agglomerative clustering. Here, singleton clusters were allowed.

#### Population analysis of channel model parameters

As a single channel model could be shared across multiple models, we mapped channel density values into the same conductance space for direct comparison. First, the analysis was constrained to parameters associated with the somatic compartment of cell models for easy comparison. Next, the “conductance density” values for each channel model within the database was extracted from NeuroML cell model files using the ElementTree XML API in the Python Standard Library. If a given channel model was not present in the cell model file, then a value of zero was set for the conductance density. If a channel model was present in the cell model file, the conductance density was expressed in units mS/cm^2^ for consistency. Thus, conductance density values across all channel models for each cell model were expressed in matrix form. This conductance density matrix was then scaled for each column vector of channel model conductance density values across the population of cell models sharing the same channel model. The maximum value was found across the cell models and the column vector divided by that maximum value. This preserved the statistical structure across cell models for the same channel model. Finally, the scaled conductance density values were then integrated to form a cumulative conductance density for the channel model cluster in the case that multiple channel models belonged to the same discovered channel model cluster and were embedded in the same cell model. This final non-negative value was referred to as the *normalized channel density* (NCD).

For the joint visualization, we restricted the visualization of the electrophysiological property space to those models with Z-score less than three within the 3-D property space. In the following statistical analysis, all models were used for computing the average NCD, as well as the bootstrapped 95% confidence intervals reported.

## Acknowledgements

We would like to acknowledge Angus Silver and all of the current and previous NeuroML Editors for NeuroML language development (https://docs.neuroml.org/NeuroMLOrg/Board). We thank the model translators who converted published models to NeuroML format (https://github.com/orgs/OpenSourceBrain/people), and those who contributed to the NeuroML toolchain (https://github.com/orgs/NeuroML/people). We also thank Charly McCown, Ashwin Rajadesingan, Harsha Velugoti Penchala, and Veer Addepalli for their contributions to NeuroML-DB.

## Financial Disclosure Statement

This work was funded in part by the National Institute on Deafness and Other Communication Disorders through award F31DC016811 to JB, by the National Institute of Mental Health through award R01MH106674 to SMC, and by the National Institute of Biomedical Imaging and Bioengineering and National Institute of Neurological Disorders and Stroke through awards R01EB021711 and U19NS112953 to RCG. VH was funded in part by the National Institute on Deafness and Other Communication Disorders through award R01DC019278. These funding institutions had no role in the study design, data collection and analysis, decision to publish, or preparation of the manuscript.

## Notes

### Competing Interest Statement

The authors have declared no competing interest.

